# Palaeogenomics reveal a hybrid origin of the world’s largest *Camelus* species

**DOI:** 10.1101/2021.10.14.464381

**Authors:** Junxia Yuan, Michael V. Westbury, Shungang Chen, Jiaming Hu, Fengli Zhang, Siren Wang, Zhen Zhang, Linying Wang, Bo Xiao, Xindong Hou, Fuqiang Li, Xulong Lai, Wenhui Liu, Guilian Sheng

## Abstract

The extinct *Camelus knoblochi* is known as the largest camel in genus *Camelus*, but its relationship to modern *Camelus* species remains unclear. In this study, we reported the first mitochondrial and nuclear analyses of seven Late Pleistocene *C. knoblochi* samples from Northeastern China. We found that they are inseparable to wild Bactrian camel on the matrilineal side, but belong to a distinct cluster on the biparental nuclear side. Further admixture proportion analyses suggested hybrid ancestry between both the ancestors of the modern wild and domesticated Bactrian camels, with ~65% contribution from the former and ~35% from the later. By calculating the coalescence time for three *Camelus* species above, we estimated the hybridization event occurred between approximately 0.8 and 0.33 Ma. We also used Bayesian skyline to reconstruct the maternal demographic trajectories for different *Camelus* to better compare their evolutionary histories. Our results provide molecular insights into *C. knoblochi* and fill in a vital piece in understanding the genus *Camelus*.

## Introduction

The genus *Camelus* contains three extant species, i.e., the one-humped Arabian camel (*C. dromedarius*), and the two-humped wild Bactrian (*C. ferus*) and domesticated Bactrian (*C. bactrianus*) camels. They are all well-adapted to arid climates, with the Arabian camel being distributed in low-altitude torrid regions across northern Africa and southwestern Asia, and the two Bactrian species inhabiting temperate dry environments from China to middle Asia (Xue et al., 2015; Mohandesan et al., 2017; Tibary and El Allali, 2020). Among extant members of *Camelus*, the wild Bactrian camel is the only feral species, while the Arabian and domesticated Bactrian camels, which have played important roles in the progress of human civilization, retain no natural populations (Qi, 2004; Almathen et al., 2016; Orlando, 2016). The genus *Camelus* also contains several fossil species, e.g., *C. knoblochi*, *C. grattatdi*, *C. sivalensis* and *C. thomasi*, among which *C. knoblochi* had the largest body size (Titov, 2008).

Bactrian camels once ranged from the great bend of the Yellow River westward to central Kazakhstan (Tulgat and Schaller, 1992), but their present wild distribution has greatly reduced in recent years (Tulgat and Schaller, 1992). Currently, the wild Bactrian camel is restricted to four isolated regions, three in China and one in Mongolia (Silbermayr et al., 2009). Although the two Bactrian species are physically isolated now, their genetic relationship has been debated for many years. There were two main hypotheses: the wild Bactrian camel was the ancestor of the domesticated, or it was feral descendant from the later (Peters and van den Driesch, 1997). Moreover, the wild Bactrian camel was once even considered as a subspecies of the domesticated one, namely *C. bactrianus ferus*. In recent years, mitochondrial and nuclear evidence suggested their clear genetic differentiation without any direct ancestral relationships (Ji et al., 2009; Silbermayr et al., 2009; Zhang et al., 2015; Ming et al., 2020), leaving a suspension for identifying their possibly respective or shared ancestors.

Based on fossil evidence, *C. knoblochi* ranged from the Sea of Azov Region to Western Transbaikalia during 0.45-0.12 million years ago (Ma). It subsequently shifted its habitat eastward at the beginning of the Late Pleistocene (approximately 0.12 Ma) and was restricted from the Urals to Northeastern China during the Late Pleistocene (Titov, 2008). It is generally thought that *C. knoblochi* became extinct at the end of Pleistocene and was ultimately replaced by Bactrian camels owing to climatic aridization and plant community transitions (Titov, 2008). Morphologically, *C. knoblochi* differs from Bactrian camels in its larger body size and broader infraorbital shelf (Geraads et al., 2019), with most other skeletal characteristics being similar. Geraads et al. (2019, 2020) suggested that *C. knoblochi* was more closely related to Arabian camels than to Bactrian camels. In contrast, another study proposed *C. knoblochi* was similar to, and potentially ancestral to Bactrian camels (Rowan et al., 2018). Therefore, the relationship between *C. knoblochi* and the extant *Camelus* is not yet well understood.

Modern and ancient DNA (aDNA) studies have deeply advanced our understanding on the phylogeny and domestication of camelid species (Cui et al., 2007; Ji et al., 2009; Wu et al., 2014; Heintzman et al., 2015; Almathen et al., 2016; Mohandesan et al., 2017; Chen et al., 2019; Lado et al., 2020; Ming et al., 2020; Díaz-Maroto et al., 2021). However, to our knowledge, no molecular analysis has been carried out on *C. knoblochi*. In this study, we retrieved palaeogenomic data from seven Late Pleistocene *C. knoblochi* remains. We explored the phylogenetic relationship between this extinct species and extant Old World camels, inferred their respective maternal demographic histories, and estimated the divergence times and genetic diversities of each camelid lineages. The new findings presented here provide us further insight into the genetic components and evolutionary history of genus *Camelus*.

## Results

### Mitochondrial and nuclear genome mapping

Seven *C. knoblochi* remains excavated from Northeastern China are included in this study, dating to between around 50 and 25.5 thousand years ago (ka) (Figure 1A; Appendix 1-table 1). Using aDNA methodology and High-throughput sequencing, we successfully obtained near-complete mitochondrial genomes from all individuals with sequencing depths ranging from 6.5 to 17.1 folds. Higher coverages were generated when mapping against the mitogenome of the wild Bactrian camel than to that of the domesticated Bactrian camel (Appendix 1-table 2). The DNA fragments are short (average length: 48-66 bp) and show evident damage characteristics (Appendix 2-figures 1-3), verifying the obtained sequences are not from modern contamination. Endogenous nuclear DNA content ranges between 0.067-2.983% and allowed the recovery of partial nuclear genomes for each sample (0.003-0.069× coverage). Detailed data processing information is listed in Appendix 1-table 2.

**Figure 1.**
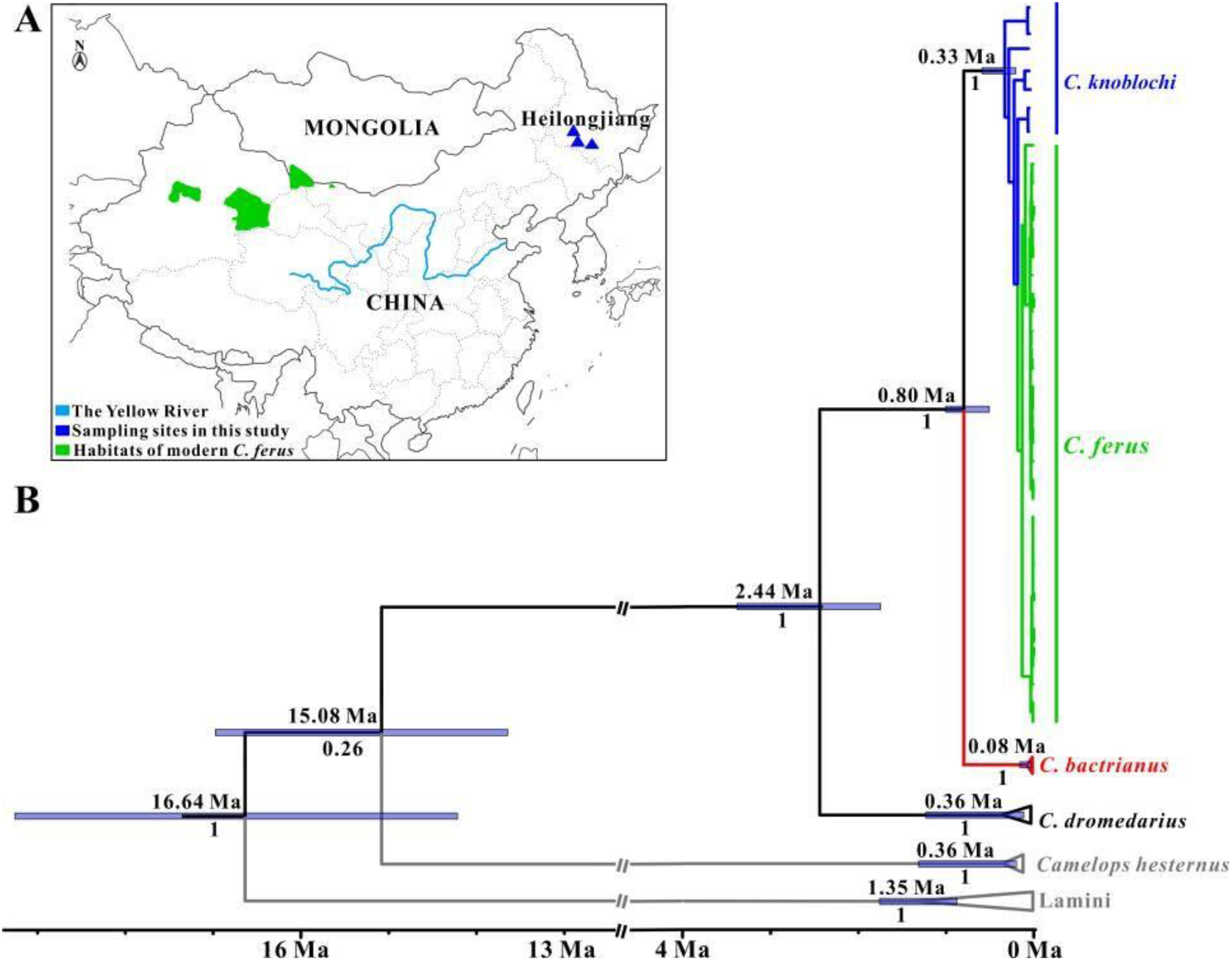
(A) Sampling locations of *C. knoblochi* analyzed in this study. (B) Maximum clade credibility tree of family Camelidae based on near-complete mitogenomes computed using BEAST. Node heights are centered on the median posterior age estimates with blue bars showing 95% credibility intervals of the divergence times. Numbers below nodes represent the posterior values. The median radiocarbon ages based on the ^14^C dating results or the median age from BEAST estimation were used as tip dates of *C. knoblochi* specimens, shown in Appendix 1-table 1. The ages of the remaining samples included in this analysis are listed in Appendix 1-table 3.

### Sex determination

We found the ratios of X chromosome coverage to autosome coverage to range from 0.52 to 0.96. While no individuals displayed the perfect ratios of 0.5 for male or one for female, all individuals were only slightly deviated from these ratios. Therefore, we could clearly see that our *C. knoblochi* dataset contained three males and four females (Appendix 1-table 2).

### Analysis of maternal phylogeny

A mitogenome BEAST phylogenetic analysis revealed that camelid individuals were divided into two clades (Figure 1B), i.e., Lamini clade and *Camelops*-*Camelus* clade, consistent with results suggested by Heintzman et al. (2015). In the *Camelops*-*Camelus* clade, *Camelops* specimens from North America formed a separate subclade while the Old World *Camelus* individuals formed another. The latter branch was further divided into three lineages, with the two species of Bactrian camels being more closely related to each other than to the Arabian camel, also consistent with previous studies (Heintzman et al., 2015; Chen et al., 2019; Ming et al., 2020). Our mitogenome BEAST tree revealed that the extinct *C. knoblochi* was nested within the diversity of extant wild Bactrian camel from Mongolia with a high posterior clade probability of 1 (Figure 1B). The entire topology is further robustly support by additional maximum likelihood (ML) tree using RAxML software (Appendix 2-figure 4).

Using the fossil calibrated node from Camelini vs. Lamini and tip calibrations, we estimated the substitution rate around 6.55×10^−9^ substitutions/site/year (95% CI: 5.26-7.92×10^−9^), which is not contradictory to previous calculations, e.g., 8.0×10^−9^ in Chen et al. (2019), and 5.0×10^−9^ in Fitak et al. (2016). The split between Old World *Camelus* and New World *Camelops* occurred during the Miocene (15.08 Ma, 95% CI: 16.97-13.64 Ma). The divergence time between Arabian and Bactrian camels is approximately 2.44 Ma (95% CI: 3.38-1.74 Ma). The coalescent time of all Bactrian species is dated around 0.80 Ma (95% CI: 1.00-0.51 Ma), while the time to the most recent common ancestor (TMRCA) of our samples plus the wild Bactrian camels is 0.33 Ma (95% CI: 0.59-0.21 Ma).

### Nuclear population structure

Our principal component analysis (PCA) using several representatives per *Camelus* species (Figure 2A) showed Arabian camel to separate first on the PC1 axis (58.9% of the variation in the data explained by PC1), and the remaining three species form three distinct clusters along the PC2 axis (7.3% of the variation). We found a near identical result when simulating aDNA damage and down sampling two individuals per extant species, suggesting that this result is biologically driven rather than by biases caused by the low coverage and damaged nature of our *C. knoblochi* data (Appendix 2-figure 5). The PCA computed using the dataset excluding Arabian camel also showed strong separation between all three species along the PC1 axis. On the PC2 axis, substructure appears within domesticated Bactrian camel. As PCs 2, 3, and 4 were explained by similar levels of variance (6.1%, 5.0%, and 4.4% respectively), we also investigated variation along these axes. Similar to PC2, PCs 3 and 4 also arise from variation within domesticated Bactrian camel (Appendix 2-figure 6).

**Figure 2.**
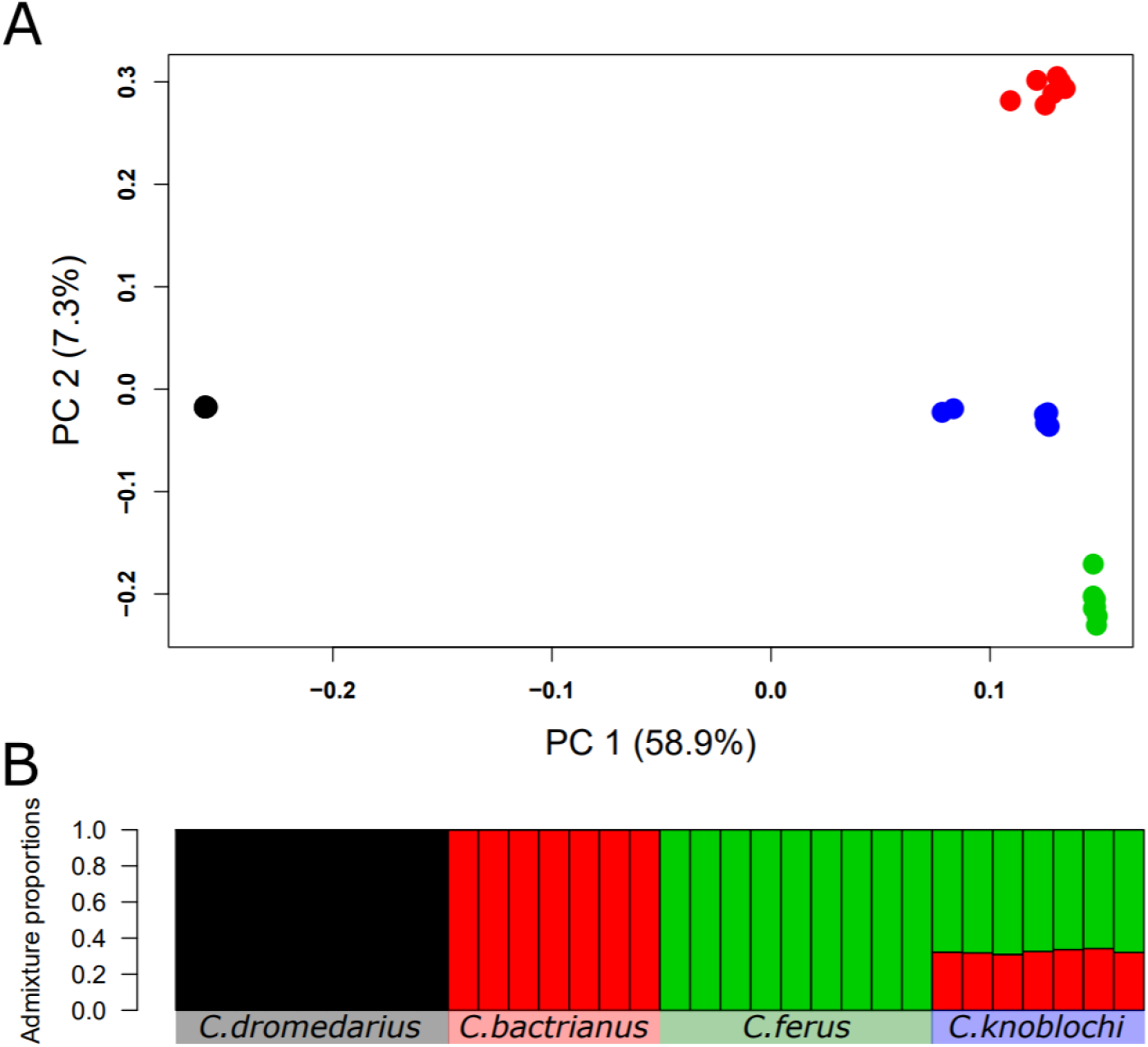
Nuclear population structure comparisons of four *Camelus* species. (A) Principal component analysis (PCA) computed using genotype likelihoods and representatives from all *Camelus* species analyzed in this study. Colours correspond to each species: black - *C. dromedarius*, red - *C. bactrianus*, green - *C. ferus*, and blue - *C. knoblochi.* (B) Admixture proportion analysis computed using genotype likelihoods and an *a priori* population number (K) of three.

Our admixture proportion analyses using all *Camelus* species converged on a K=3 but not on K=4, suggesting that there are three and not four populations in our dataset. This analysis showed the extant species forming their own populations with no migration between them, and *C. knoblochi* to contain ~35% domesticated Bactrian camel and ~65% wild Bactrian camel ancestry (Figure 2B). We found a similar result when using the aDNA simulated dataset (Appendix 2-figure 7). However, with the aDNA simulated dataset, K=4 did converge. With K=4, we start to uncover some substructure within the domesticated Bactrian camel while *C. knoblochi* still appears as a mixture between the ancestors of domesticated and wild Bactrian camels (Appendix 2-figure 7). Due to this convergence, we also looked into the K=4 NGSadmix output with the best likelihood for the original dataset. Although it did not converge, we see a similar pattern of substructure within the domesticated Bactrian camel arising as with the aDNA simulated dataset (Appendix 2-figure 8).

### Population dynamics

Bayesian skyline analysis of wild Bactrian camel mitogenomes indicated a steep decline in female effective population size in the last several hundred years, while the ancestor of domesticated Bactrian camels showed population expansion during the past 80 thousand years. *C. knoblochi* kept a relatively steady maternal population size during the Late Pleistocene, only observing a slight increase of 95% credibility intervals between 55 and 50 ka (Figure 3).

**Figure 3.**
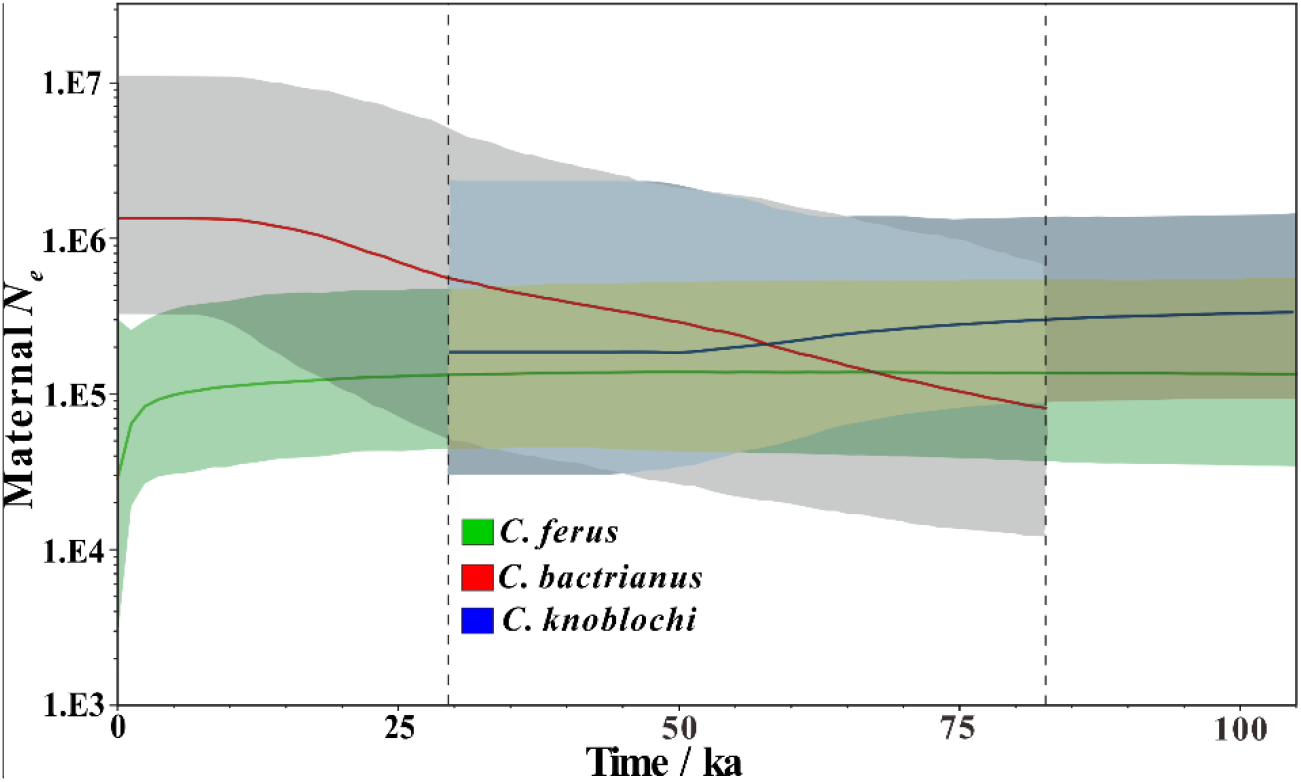
Bayesian skyline plot of the two Bactrian camels and *C. knoblochi* individuals based on near-complete mitochondrial genomes. Green, red and blue lines indicate median female *N*_*e*_ of *C. ferus*, *C. bactrianus* and *C. knoblochi* changes over time. Correspondingly, the grey, blue and green shaded areas indicate their 95% credibility intervals.

### Mitochondrial DNA Diversity

We also calculated genetic diversity of *Camelus* lineages (Table 1). The lowest genetic diversity is observed in modern wild Bactrian camel, while the Late Pleistocene *C. knoblochi* shows higher mitochondrial diversity compared to extant Old World camels.

**Table 1.** Genetic diversity of *Camelus* lineages based on mitogenomes.

| Taxon | *C. ferus* | *C. bactrianus* | *C. dromedarius* | *C. knoblochi* |
| --- | --- | --- | --- | --- |
| Genetic diversity | 0.001162 | 0.001298 | 0.001725 | 0.003002 |

## Discussion

### Phylogenetic position and origin for *Camelus knoblochi*

All fossils in this study are diagnostic and have been carefully evaluated in morphology (Appendix 2-figures 9-13). They have generally larger body sizes than extant *Camelus* materials. We confirm that size differences are not from the gender biases as our molecular sex determination suggested that both female and male individuals were included. Therefore, our samples are clearly attributed to *C. knoblochi*.

We reconstructed phylogenetic trees of camelids based on near-complete mitochondrial genomes. Both Bayesian and ML approaches showed that our seven *C. knoblochi* do not form a separate lineage but to cluster with the extant wild Bactrian camel (Figure 1B; Appendix 2-figure 4). Such intermixed pattern is also observed in other mammals, for example, polar vs. brown bear (Miller et al., 2012) and Tibetan wild ass vs. Mongolian wild ass (Vilstrup et al., 2013). These phenomenon could be explained by past hybridization events or incomplete lineage sorting, which might lead to the discrepancies between molecular and morphological identification (Taron et al., 2021). Therefore, it is imperative to also consider nuclear data as the mitochondrial marker may not reflect the full picture of the species’ evolutionary history (Miller et al., 2012; Sheng et al., 2019).

Contrary to mitochondrial results, the nuclear analyses clearly suggest that *C. knoblochi* is a unique group, which is closer to wild Bactrian camel than to domesticated Bactrian camel (Figures 2A & 1B). Using simulated aDNA dataset, we verified this result while considering potential biases linked to the low genomic coverage or DNA damage (Appendix 2-figures 5-8). Further admixture proportion analyses revealed that *C. knoblochi* shares ancestry from both wild Bactrian camels and the ancestor of modern domesticated Bactrian camels. This extinct camel is mostly a hybrid between the ancestors of two modern Bactrian camel species. The fact that our samples contain unilateral matrilineage (Figure 1B) suggests *C. knoblochi* arose from mating of wild Bactrian camel females and the male ancestors of modern domesticated Bactrian camel. However, we cannot know if this is always the case as we covered limited geographical sampling sites. Previous studies have indicated that hybridization is common in extant *Camelus* species, for instance in modern domesticated Bactrian camel × wild Bactrian camel (Felkel et al., 2019; Ming et al., 2020) and domesticated Bactrian camel × Arabian camel (Burger, 2016; Ming et al., 2020). Our analyses detect an ancient mixed ancestry, which means hybridization may also be common to Pleistocene *Camelus*. However, it is noteworthy that the earliest known remains of the two Bactrian camels were limited in the Late Pleistocene, while the earliest *C. knoblochi* fossil recorders are traced back to Middle Pleistocene (Titov, 2008). Further investigations on fossil materials from the three *Camelus* species above are needed to fully understand their evolutionary histories.

### Divergence time for *Camelus* species

The family Camelidae originated in North America approximately 45-40 Ma (Stanley et al., 1994). This was followed by the origin of the genus *Paracamelus*, most likely ancestral to all *Camelus*, which dispersed from North America into Eurasia and Africa via the Bering Strait around 6.3-5.8 Ma (van der Made et al., 2002). For all Old World camels, *Camelus grattardi* is thought to be the earliest representative based on remains from the Plio-Pleistocene strata (2.9-2.2 Ma) of Ethiopia (Geraads et al., 2019). Using root-and-tip dating calibrations in BEAST (Figure 1B), we estimated the subsequent split between Arabian and Bactrian camels around 2.44 Ma, much younger than previous molecular dating (8-4.1 Ma) (Cui et al., 2007; Wu et al., 2014; Heintzmann et al., 2015). However, Geraads et al. (2020) argued that the above mentioned problematic calibration points probably led to overestimation and newly suggested 2-1 Ma for that node using fossil data. As the fossil record tends to provide minimal age estimates (Geraad et al., 2020), we confirm our estimation to be reasonable.

The divergence between the ancestors of domesticated and wild Bactrian camels occurred around 0.8 Ma, indicating the temporal upper limit for their hybridization. The earliest *C. knoblochi* fossils are dated back to the second half of the Middle Pleistocene (0.45-0.12 Ma) (Titov, 2008) and the coalescence for all *C. knoblochi* here was traced to approximately 0.33 Ma (Figure 1B). Hence, the now extinct *C. knoblochi* likely arose due to a hybridization event occurred between 0.8 and 0.33 Ma (Figure 4). However, it is still unclear where hybridization occurred and how *C. knoblochi* became a fixed unique lineage. Genetic investigation into more Pleistocene camels should be essential to answer these questions.

**Figure 4.**
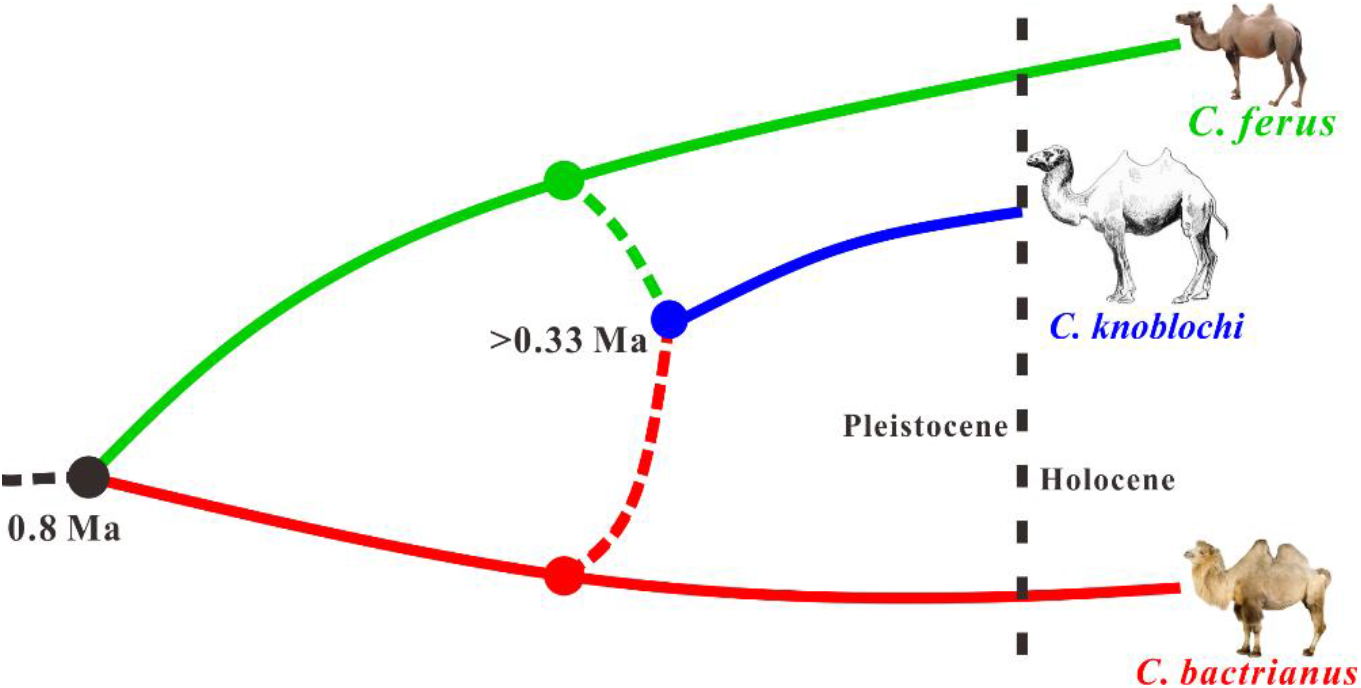
Hypothesis for evolutionary outline of *C. knoblochi* and the two Bactrian camels based on BEAST, PCA and admixture proportion analyses.

### Genetic diversities and demographic histories for *Camelus*

We investigated genetic diversity of different lineages within *Camelus* (Table 1). Similar to previous studies (Mohandesan et al., 2017; Ming et al., 2020), wild Bactrian camels display the lowest diversity value (0.001162). This is not unexpected as wild Bactrian camels are suffering from a strong loss of genetic diversity in recent years, which may be linked to geographic isolation and small population size. They are currently limited to four isolated geographic regions (Silbermayr et al., 2009) and have only ~1,000 individuals in China and Mongolia (Xue et al., 2015; Burger et al., 2019). In addition, our BSP analysis revealed that the maternal population size of wild Bactrian camels remained relatively stable since 100 ka, except a sharp decline in the last few hundred years (Figure 3), possibly owing to human activities. However, the maternal population size of domesticated Bactrian camels kept a continued growth during the past 80 ka, reflecting that the ancestors of the wild and domesticated Bactrian camels experienced different pressures and evolutionary histories during the Late Pleistocene.

The available Late Pleistocene *C. knoblochi* individuals exhibit higher genetic diversity compared to extant *Camelus* (Table 1). This may be due to its hybrid origin or might indicate a relatively larger population size during the Late Pleistocene. Fossil records suggested that *C. knoblochi* was widely distributed in temperate and cold steppes of Central and East Asia during the Late Pleistocene. Bayesian skyline reconstruction showed the maternal effective population size of *C. knoblochi* lineage remained generally stable during the Late Pleistocene, while later maintaining a large population (Figure 3). The observed expansion could correspond to the early substage of Guxiangtun periglacial stage (70-53 ka). The climate was very cold in Northeast China during this period and the average annual temperature might be 10 ℃ lower than that of today (Sun et al., 1985). After this cold stage, there is a slow population increase of *C. knoblochi*. The demographic history of *C. knoblochi* may be further explored by involving more individuals in future studies.

## Methods

### Sampling

A total of seven *Camelus* specimens were collected from three regions, i.e., Qinggang County, Zhaodong County, and Haerbin City, in Heilongjiang Province, Northeastern China (Figure 1A). Four of them (CADG387, CADG388, CADG389 and CADG596) are stored in Zhaoyuan Museum, two samples (CADG529 and CADG533) in Daqing Museum and one sample (CADG665) in Yifu Museum of China University of Geosciences. All specimens are assigned to *C. knoblochi* based on morphological characters. Detailed information of these specimens is shown in Appendix 1-table 1 and Appendix 2-figures 9-13.

### DNA extraction, library preparation and sequencing

DNA extraction and library preparation were all conducted in dedicated clean room facilities at China University of Geoscience (Wuhan). Approximately 200 mg of each specimen was ground into fine powder. DNA extraction was carried out as described previously (Hu et al., 2021). DNA was eluted twice by adding EB buffer to produce 80 μL of final DNA extract for each sample. DNA extracts were converted into double-stranded Illumina sequencing libraries following the protocol from Meyer and Kircher (2010) with several modifications (Hu et al., 2021). Finally, indexed libraries were quantified, pooled, and sequenced using an Illumina HiSeqX10 platform in a 2 × 150 bp paired-end run. Additionally, extraction and library blanks were included throughout all steps to monitor potential contamination.

### Mitochondrial data processing

Raw reads were trimmed of adapters using Cutadapt v1.4.2 (Martin, 2011), requiring a minimum overlap of 1 bp, and discarding reads shorter than 30 bp. The trimmed reads were then mapped to modified mitogenome reference sequences of wild Bactrian camel (Genbank No. EF212038) and domesticated Bactrian camel (Genbank No. EF507799), independently, using the “aln” algorithm in Burrows-Wheeler aligner (BWA) v0.6.2 (Li and Durbin, 2010) with default parameters. Due to the circular state of the mitochondrial genome, the modified references had 20 bp added to the ends from the opposite edge. Reads with a MapQuality score less than 30 and designated as PCR duplicates were removed using SAMtools v0.1.9 (Li et al., 2009). The final mitochondrial consensus sequences were generated in Geneious (https://www.geneious.com/), using a minimum coverage of 2 and a base agreement greater than 75%.

### Mitochondrial phylogenetic analysis

To investigate the phylogenetic position of *C. knoblochi*, a Bayesian phylogenetic analysis was performed in BEAST v1.8.2 (Drummond et al., 2012) using the near-complete mitochondrial genomes (Appendix 1-table 3). The dataset contains 7 *C. knoblochi*, 15 *C. bactrianus*, 32 *C. ferus*, 10 *C. dromedarius*, 2 *Lama glama*, 3 *Lama pacos*, 5 *Lama guanicoe*, 2 *Vicugna Vicugna,* 3 *Camelops* cf. *hesternus*, and one *Sus scrofa* as the outgroup. A multiple sequence alignment was generated by using MAFFT v7.471 (Katoh et al., 2002), and the problematic section of the control region was removed. The final alignment of the combined dataset contains 16,680 nucleotide positions. PartitionFinder v2.1.1 (Lanfear et al., 2016) was used to identify the optimal partitioning scheme, and our dataset was divided into eight distinct partitions. A root-and-tip-dating approach was used for the calibrated analysis. Firstly, based on the first appearance of Lamini, the divergence time between Camelini and Lamini was set at 17.5 Ma with a standard deviation of 1.52 Ma. Secondly, tip dates were considered zero for the modern individuals and the median radiocarbon or stratigraphic ages for the ancient specimens (Appendix 1-tables 1, 3). As to the ancient specimens with no stratigraphic information and also beyond the limits of radiocarbon dating, the median age from BEAST estimation was used as the sample date (Appendix 1-table 1). The BEAST analysis was conducted under a relaxed uncorrelated lognormal molecular clock. A coalescent Bayesian skyline tree model and the GTR+G substitution model were selected. Markov chain Monte Carlo chains were performed with 100,000,000 generations and sampled every 10,000 states. The results were analyzed using TRACER v1.7 (Rambaut et al., 2018) to check the convergence of each parameter and sufficient sampling (ESS>200). The first 25,000,000 of trees were discarded as burn-in, the Maximum Clade Credibility tree was summarized using the TreeAnnotator v1.5.4 (Drummond et al., 2012), which was visualized using FigTree v1.4.4 (http://tree.bio.ed.ac.uk/software/figtree). Moreover, Bayesian skyline analyses were conducted to infer female effective population size changes over time in the wild Bactrian camels, the ancestor of the modern domesticated Bactrian camels, and *C. knoblochi* lineages using BEAST v1.8.2 (Drummond et al., 2012).

Genetic diversity of *C. knoblochi* and extant *Camelus* individuals (Appendix 1-table 3) was calculated using MEGA 7 software (Kumar et al., 2016), all positions containing gaps or missing data were excluded and the final dataset had a length of 12,285 bp.

### Nuclear data processing

For the ancient *C. knoblochi* samples, we followed the same mapping protocol as for the mitochondrial genomes, except we used the wild Bactrian camel nuclear genome (GCF_009834535.1) instead of the mitochondrial genome as reference. As a comparative dataset, we downloaded the raw sequencing reads from nine Arabian camel, seven domesticated Bactrian camel, and nine wild Bactrian camel (Fitak et al., 2020). We merged overlapping paired end reads using FLASH v1.2.11 (Magoč and Salzberg, 2011) utilizing default parameters. We mapped both merged and unmerged reads to the available wild Bactrian camel reference nuclear genome using BWA v0.7.15 (Li and Durbin, 2010) utilizing the mem algorithm and otherwise default parameters. We parsed the alignment files and removed duplicates and reads of mapping quality score <30 using SAMtools v1.6 (Li et al., 2009). Mapping statistics can be found in Appendix 1-tables 2, 4.

### Sex determination

We determined the sex of our *C. knoblochi* individuals by comparing the average coverage of reads mapping to the X chromosome, to that of a chromosome of similar length (chromosome 1). We did this using SAMtools depth and either specifying the X chromosome or chromosome 1 using the parameter -b, and to include sites with no coverage (-a). This method works as males should have half the amount of coverage on the X chromosome compared to the autosomes due to only having a single copy of the X chromosome but two copies of each autosome, whereas females should have relatively similar coverages for both the X chromosome and autosomes.

### Nuclear population structure

We evaluated the nuclear relationships between the four *Camelus* species included in this study by performing a PCA, as well as estimating admixture proportions. We performed these analyses twice, once using all four species, and once excluding Arabian camel. Due to the very low coverage manner of our *C. knoblochi* individuals, we performed these analyses using genotype likelihoods (GL) computed with ANGSD v0.921 (Korneliussen et al., 2014). We computed GL for all individuals specifying the parameters: minimum mapping and base qualities of 30 (-minmapQ 30 -minQ 30), calculate genotype likelihoods using the GATK algorithm (-GL 2), output a beagle GL file (-doGlf 2), calculate major and minor alleles based on GL (-doMajorMinor 1), remove transitions (-rmtrans 1), only include SNPs with a p-values <1e-6 (-SNP_pval 1e-6), only consider autosomal chromosomes (-rf), a minimum minor allele frequency of 0.05 (-minmaf 0.05), skip triallelic sites (-skiptriallelic 1), only consider reads mapping to one region uniquely (-uniqueonly 1), only consider sites if at least 26 individuals have coverage (-minind 26). When calculating GL but excluding Arabian camel we used the same parameters but only considered sites if at least 17 individuals had coverage. To construct a covariance matrix for PCA we used PCAngsd v0.98 (Meisner and Albrechtsen, 2018). To estimate ancestry components we used NGSadmix v32 (Skotte et al., 2013) specifying the number of *a priori* populations (K) as 3 or 4. To evaluate the reliability of our NGSadmix results, we repeated the NGSadmix analysis up to 100 times, printing the likelihood each time. If the best likelihood was obtained twice, the results were considered to have converged and were therefore reliable. If this was not obtained within 100 replicates, then the reliability of the results could not be determined.

### Palaeogenomic data reliability assessment

Due to the large disparity between data quality and coverage of the modern and ancient datasets, we assessed whether our results could be data driven as opposed to biological. To test this we randomly selected two individuals from each of the extant species, reduced their raw read lengths down to 40 bp, and added aDNA damage patterns using TAPAS v1.2 (Taron et al., 2018) and the mapdamage output from our *C. knoblochi* individual with the highest coverage (CADG533). We mapped the artificially damaged reads to the wild Bactrian camel reference genome as was done with the empirical ancient data and subsampled the resultant bam files to ~0.07x with SAMtools. We reran the PCA and admixture proportion analyses with the aDNA simulated data for these individuals instead of the original high quality data. To ensure comparability, we used the same sites as we had recovered with the complete dataset.

## Acknowledgements

We are grateful to Mr. Hailong Ji (Paleontological fossil conservation center of Qinggang County) for his help in collecting the samples. This work was supported by the National Natural Science Foundation of China (grant nos. 41472014, 42172027).

## Author contributions

Junxia Yuan, Xulong Lai and Guilian Sheng conceived the study; Shungang Chen, Linying Wang, Jiaming Hu, Bo Xiao and Xindong Hou performed the experiments; Junxia Yuan, Guilian Sheng and Michael V. Westbury guided the experiment and bioinformatics analyses; Shungang Chen, Michael V. Westbury, Linying Wang and Junxia Yuan analyzed the data; Fengli Zhang, Siren Wang, Zhen Zhang and Fuqiang Li helped to collect the samples; Wenhui Liu carried out morphological analyses of the samples; Junxia Yuan, Michael V. Westbury, Guilian Sheng, Wenhui Liu, Jiaming Hu, Shungang Chen, Linying Wang and Xulong Lai wrote the paper. All authors read and gave comments to the final version of the manuscript.

## Data availability

The obtained mitochondrial genomes reported in this article have been deposited in the Genbank under the accession nos. MZ430515-MZ430521.

## Supplementary Materials

**Appendix 1-table 1.** The information of the *C. knoblochi* specimens analyzed in this study.

**Appendix 1-table 2.** Raw reads mapping results of the *C. knoblochi* specimens analyzed in this study.

**Appendix 1-table 3.** Sequences downloaded from NCBI for mitochondrial data analyses in this study.

**Appendix 1-table 4.** Raw reads mapping statistic results of modern Old World camels download from NCBI.

**Appendix 2-figure 1.** Cytosine deamination frequency inferred from *C. knoblochi* samples analyzed in this study.

**Appendix 2-figure 2.** Estimated endogenous fragment length distributions for *C. knoblochi* samples analyzed in this study.

**Appendix 2-figure 3.** Coverage plots of the mitochondrial genomes obtained in this study.

**Appendix 2-figure 4.** Maximum-likelihood phylogenetic tree of camelids based on complete mitochondrial genomes.

**Appendix 2-figure 5.** Principal component analysis computed using genotype likelihoods with *C. dromedarius*.

**Appendix 2-figure 6.** Principal component analysis computed using genotype likelihoods without *C. dromedarius*.

**Appendix 2-figure 7.** Admixture proportions analyses computed using genotype likelihoods and an *a priori* population numbers (K) of three and four.

**Appendix 2-figure 8.** Admixture proportions analysis computed using genotype likelihoods and an *a priori* population number (K) of four with the complete dataset.

**Appendix 2-figure 9.** The axial skeleton of *C. knoblochi* from Northeastern China.

**Appendix 2-figure 10.** The complete forearm of *C. knoblochi* from Northeastern China.

**Appendix 2-figure 11.** The incomplete forearm of *C. knoblochi* from Northeastern China.

**Appendix 2-figure 12.** The incomplete femur of *C. knoblochi* from Northeastern China.

**Appendix 2-figure 13.** The metatarsus of *C.knoblochi* from Northeastern China.

