## Supplemental materials for "Palaeogenomics reveal a hybrid origin of the world’s largest *Camelus* species"

### Supplementary Materials

#### Appendix 1

**Appendix 1-table 1. The information of the *C. knoblochi* specimens analyzed in this study.**

| Lab No. | Museum No. | Skeletal element | Location | Radiocarbon lab No. | Radiocarbon age ($^{14}C$, BP) | Calibrated radiocarbon age (BP) |
|---|---|---|---|---|---|---|
| CADG387 | ZYB:3145 | 4th cervical vertebra | Qinggang | Beta-543610 | 25,580±100 | 95.4% (30,148-29,370) |
| CADG388 | ZYB:3144 | metatarsus | Qinggang | Beta-539398 | 33,940±230 | 95.4% (38,956-37,750) |
| CADG389 | ZYB:3147 | femur | Haerbin | Beta-543611 | 39,220±400 | 95.4% (43,696-42,413) |
| CADG529* | H35364 | mandible | Qinggang | Beta-549493 | >43,500 | NA |
| CADG533 | H33979 | Radioulnare | Zhaodong | Beta-549494 | 38,830±440 | 95.4% (43,385-42,132) |
| CADG596** | ZYB:3146 | Radioulnare | Qinggang | Beta-561363 | >43,500 | NA |
| CADG665 | x0221-01 | metatarsus | Qinggang | Beta-549496 | 40,480±430 | 95.4% (44,846-43,223) |

* The median age from BEAST estimation is 48,521 BP.

** The median age from BEAST estimation is 50,083 BP.

**Appendix 1-table 2. Raw reads mapping results of the *C. knoblochi* specimens analyzed in this study.**

| | Whole genome reference | | | | | | Mitogenome reference | | | | |
|---|---|---|---|---|---|---|---|---|---|---|---|
| | GCF_009834535.1 (*C. ferus*) | | | | | | EF212038 (*C. ferus*) | | | EF507799 (*C. bactrianus*) | |
| Lab No. | Endogenous DNA (%) | Coverage (Average) | Coverage (Chromosome 1) | Coverage (X) | Ratio (X/Chromosome 1) | Sex | Sequence length (bp) | Average depth | GenBank No. | Sequence length (bp) | Average depth |
| CADG387 | 0.828 | 0.025 | 0.0250 | 0.0231 | 0.92 | Female | 16,459 | 8.8 | MZ430515 | 16,030 | 7.9 |
| CADG388 | 0.632 | 0.037 | 0.0377 | 0.0195 | 0.52 | Male | 16,245 | 8.5 | MZ430516 | 15,743 | 7.6 |
| CADG389 | 0.067 | 0.003 | 0.00314 | 0.00286 | 0.91 | Female | 14,699 | 6.5 | MZ430517 | 13,984 | 5.8 |
| CADG529 | 0.141 | 0.004 | 0.00360 | 0.00346 | 0.96 | Female | 16,611 | 14.3 | MZ430518 | 16,413 | 12.5 |
| CADG533 | 2.983 | 0.069 | 0.0769 | 0.0403 | 0.52 | Male | 16,590 | 15.6 | MZ430519 | 16,381 | 13.9 |
| CADG596 | 0.649 | 0.037 | 0.0379 | 0.0348 | 0.92 | Female | 16,433 | 17.1 | MZ430520 | 16,311 | 15.6 |
| CADG665 | 0.511 | 0.027 | 0.0262 | 0.0136 | 0.52 | Male | 15,949 | 7.5 | MZ430521 | 15,232 | 6.5 |

**Appendix 1-table 3. Sequences downloaded from NCBI for mitochondrial data analyses in this study*.**

| genus | Taxon | NCBI Accession |
|---|---|---|
| *Camelus* | *C. dromedarius* | EU159113, JN632608_(0), KU605059_(1450), KU605061_(592), KU605071_(1375), KU605072, KU605073_(0), KU605074_(0), KU605075, KU605076, KU605077, KU605078_(0), KU605079_(0), KU605080, KX554931_(0), KX554932, KX554933_(0), KX554934, NC_009849 |
| | *C. bactrianus* | AP003423, EF212037, EF507798, EF507799, KU666460, KU666461, KU666462, KU666463, KU666464, KU666465, KX554925, KX554926, KX554927, KX554928, KX554929, KX554930, MH109872_(0), MH109873, MH109874, MH109875, MH109876, MH109877, MH109878, MH109879_(0), MH109880, MH109881_(0), MH109882, MH109883_(0), MH109884, MH109885, MH109886, MH109887, MH109888, MH109889, MH109890, MH109891, MH109892, MH109893, MH109894, MH109895, MH109896, MH109897, MH109898, MH109899, MH109900_(0), MH109901, MH109902, MH109903, MH109904, MH109905, MH109906_(0), MH109907, MH109908, MH109909_(0), MH109910_(0), MH109911, MH109931, MH109932, MH109933, MH109934, MH109935, MH109936, MH109937, MH109938, MH109939, MH109940, MH109941, MH109942_(0), MH109943, MH109944, MH109945, MH109946, MH109947, MH109948, MH109949, MH109950, MH109951, MH109952, MH109953, MH109954, MH109955, MH109956, MH109957_(0), MH109958, MH109959, MH109960, MH109961, MH109962_(0), MH109963, MH109964, MH109965, MH109966, MH109967, MH109968, MH109969, MH109970, MH109971, MH109972, MH109973, MH109975, MH109976, MH109977, MH109978, MH109979, MH109980, MH109981_(0), MH109982, MH109983, MH109984, MH109986, MH109987, MH109988, MH109989, MH109990_(0), MH109992, MH109993, MH109994, MH109995, MH109996_(0), MH109997, MW357598_(1235), NC_009628 |
| | *C. ferus* | EF212038_(0), EF507800_(0), EF507801_(0), KU666451_(0), KU666452_(0), KU666453_(0), KU666454_(0), KU666455_(0), KU666456_(0), KU666457_(0), KU666458_(0), KU666459_(0), MH109912_(0), MH109913_(0), MH109914_(0), MH109915_(0), MH109916_(0), MH109917_(0), MH109918_(0), MH109919_(0), MH109920_(0), MH109921_(0), MH109922_(0), MH109923_(0), MH109924_(0), MH109925_(0), MH109926_(0), MH109927_(0), MH109928_(0), MH109929_(0), MH109930_(0), NC_009629_(0) |

**Continued Appendix 1-table 3.**

| genus | Taxon | NCBI Accession |
|---|---|---|
| *Camelops* | *C.* cf. *hesternus* | KR822420_ (102500), KR822421_(102500), KR822422_(102500) |
| *Lama* | *L. glama* | AP003426_(0), NC_012102_(0) |
| | *L. guanicoe* | EU681954_(0), KX388532_(417), KX388533_(417), KX388534_(417), NC_011822_(0) |
| *Vicugna* | *V. pacos* | AJ566364_(0), NC_002504_(0), Y19184_(0) |
| | *V. Vicugna* | FJ456892_(0), NC_013558_(0) |
| *Sus* | *S. scrofa* | KT279758_(0) |

*Ages used in BEAST analysis are indicated in parentheses following the accession number.

**Appendix 1-table 4. Raw reads mapping statistic results of modern Old World camels download from NCBI.**

| SRA accession | Species | Unmerged reads mapping | Merged reads mapping | Genome-wide Coverage | Chr1 coverage | X coverage | X:Chr1 Ratio | Sex |
|---|---|---|---|---|---|---|---|---|
| SRR1947230 | *C. dromedarius* | 313,234,390 | 1,296,312 | 15.04 | 14.68 | 14.41 | 0.98 | Female |
| SRR1947231 | *C. dromedarius* | 256,206,365 | 1,207,137 | 12.28 | 12.82 | 12.54 | 0.98 | Female |
| SRR1947232 | *C. dromedarius* | 310,856,710 | 966,452 | 14.91 | 15.17 | 14.78 | 0.97 | Female |
| SRR1947233 | *C. dromedarius* | 299,684,310 | 935,801 | 14.37 | 14.55 | 14.20 | 0.98 | Female |
| SRR1947234 | *C. dromedarius* | 307,703,032 | 1,383,181 | 14.76 | 15.33 | 14.90 | 0.97 | Female |
| SRR1947235 | *C. dromedarius* | 314,638,458 | 1,424,027 | 15.12 | 15.39 | 14.76 | 0.96 | Female |
| SRR1947236 | *C. dromedarius* | 207,474,001 | 751,729 | 9.95 | 11.35 | 5.98 | 0.53 | Male |
| SRR1947237 | *C. dromedarius* | 204,263,125 | 758,557 | 9.80 | 10.72 | 5.75 | 0.54 | Male |
| SRR1947238 | *C. dromedarius* | 221,078,741 | 990,739 | 10.62 | 10.98 | 10.63 | 0.97 | Female |
| SRR1947239 | *C. bactrianus* | 309,192,461 | 1,099,672 | 14.82 | 15.23 | 14.69 | 0.96 | Female |
| SRR1947240 | *C. bactrianus* | 312,544,529 | 1,043,500 | 14.96 | 15.44 | 8.38 | 0.54 | Male |
| SRR1947241 | *C. bactrianus* | 296,494,837 | 1,227,675 | 14.23 | 14.53 | 7.82 | 0.54 | Male |
| SRR1947242 | *C. bactrianus* | 311,793,924 | 1,514,738 | 14.99 | 14.87 | 14.35 | 0.96 | Female |
| SRR1947243 | *C. bactrianus* | 323,362,387 | 1,415,191 | 15.53 | 16.18 | 15.30 | 0.95 | Female |
| SRR1947244 | *C. bactrianus* | 322,869,734 | 1,273,280 | 15.50 | 15.33 | 14.82 | 0.97 | Female |
| SRR1947245 | *C. bactrianus* | 313,894,548 | 1,134,661 | 15.05 | 15.26 | 8.27 | 0.54 | Male |
| SRR1947246 | *C. ferus* | 292,829,738 | 1,402,101 | 14.06 | 14.16 | 13.70 | 0.97 | Female |
| SRR1947247 | *C. ferus* | 319,653,657 | 2,115,916 | 15.39 | 15.79 | 8.49 | 0.54 | Male |
| SRR1947248 | *C. ferus* | 315,142,273 | 1,177,289 | 15.12 | 15.57 | 8.35 | 0.54 | Male |
| SRR1947249 | *C. ferus* | 314,029,906 | 1,017,580 | 15.05 | 15.17 | 8.20 | 0.54 | Male |
| SRR1947250 | *C. ferus* | 333,149,533 | 1,911,141 | 16.02 | 16.01 | 15.66 | 0.98 | Female |
| SRR1947251 | *C. ferus* | 313,880,897 | 1,221,034 | 15.07 | 14.47 | 14.20 | 0.98 | Female |
| SRR1947252 | *C. ferus* | 311,897,973 | 1,171,132 | 14.95 | 15.21 | 8.23 | 0.54 | Male |
| SRR1947253 | *C. ferus* | 326,691,884 | 1,293,504 | 15.68 | 15.60 | 14.98 | 0.96 | Female |
| SRR1947254 | *C. ferus* | 312,462,764 | 1,073,956 | 14.98 | 15.13 | 8.22 | 0.54 | Male |

### Appendix 2

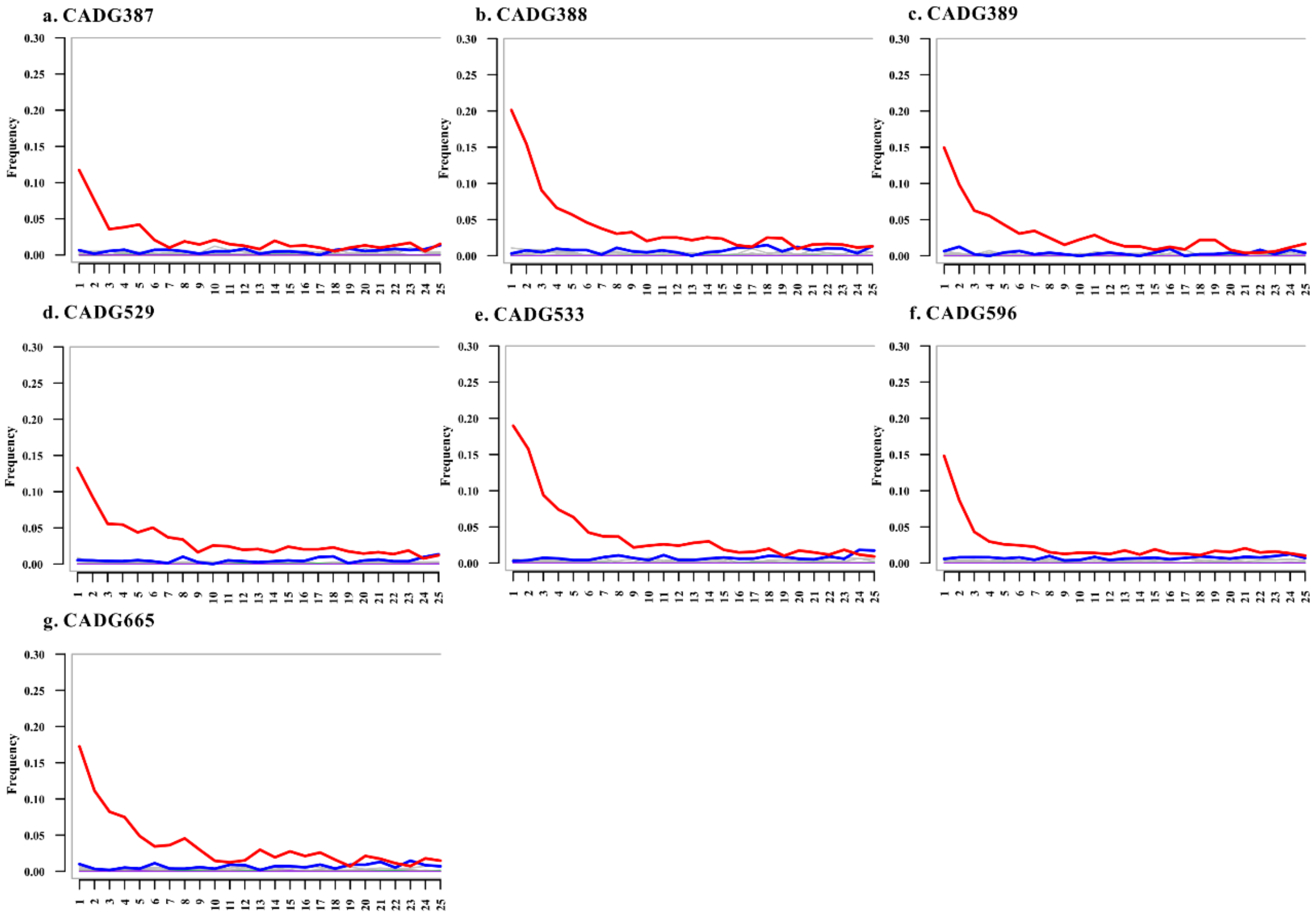


**Appendix 2-figure 1. Cytosine deamination frequency inferred from *C. knoblochi* samples analyzed in this study.** Red lines show the rates of C to T substitutions for the first 25 bases of the 5' end of the fragments that could be mapped to the *C. ferus* mitochondrial genome.

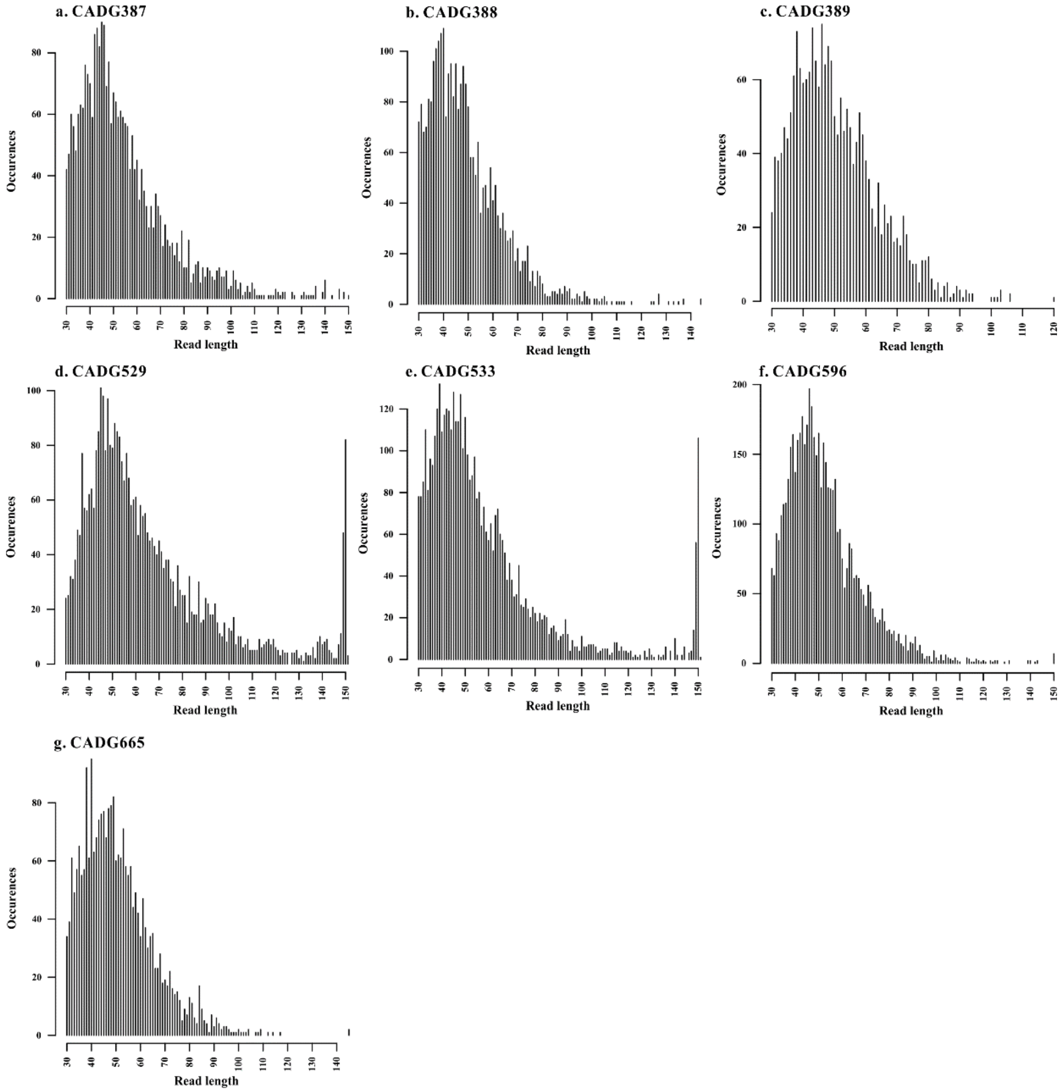


**Appendix 2-figure 2. Estimated endogenous fragment length distributions for *C. knoblochi* samples analyzed in this study.**

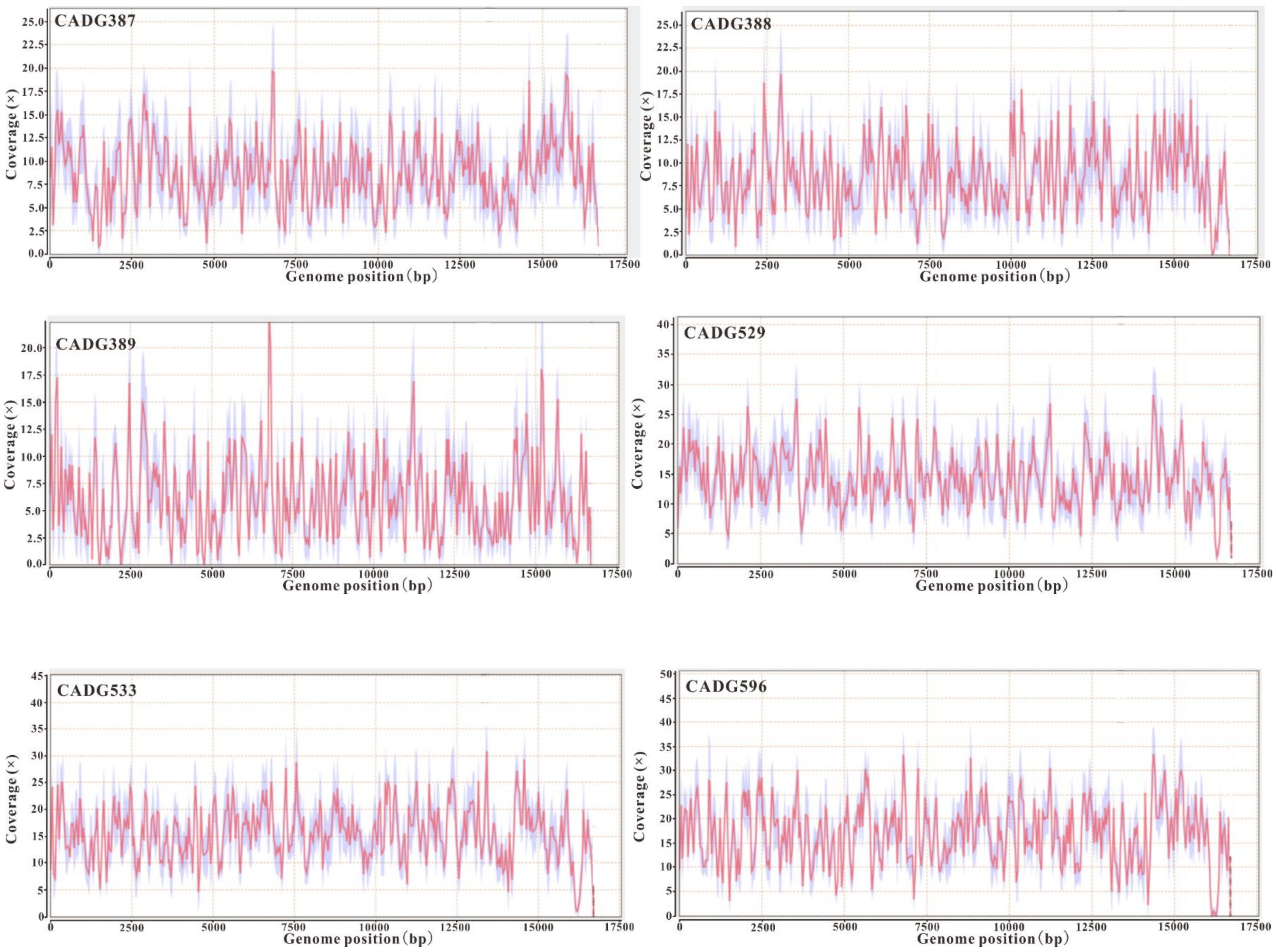
CADG387
CADG388
CADG389
CADG529
CADG533
CADG596
Coverage (×)
Genome position (bp)

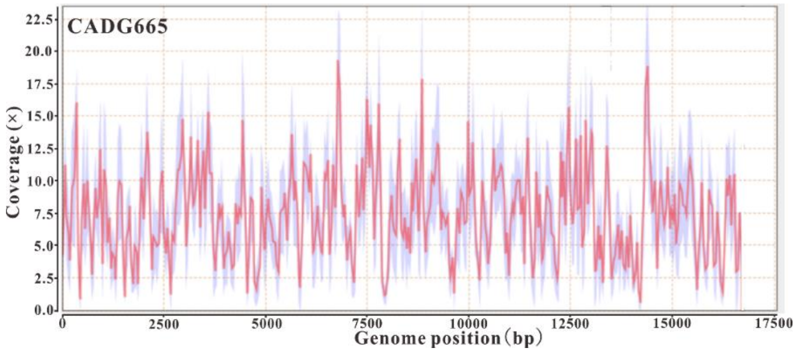


**Appendix 2-figure 3. Coverage plots of the mitochondrial genomes obtained in this study.**

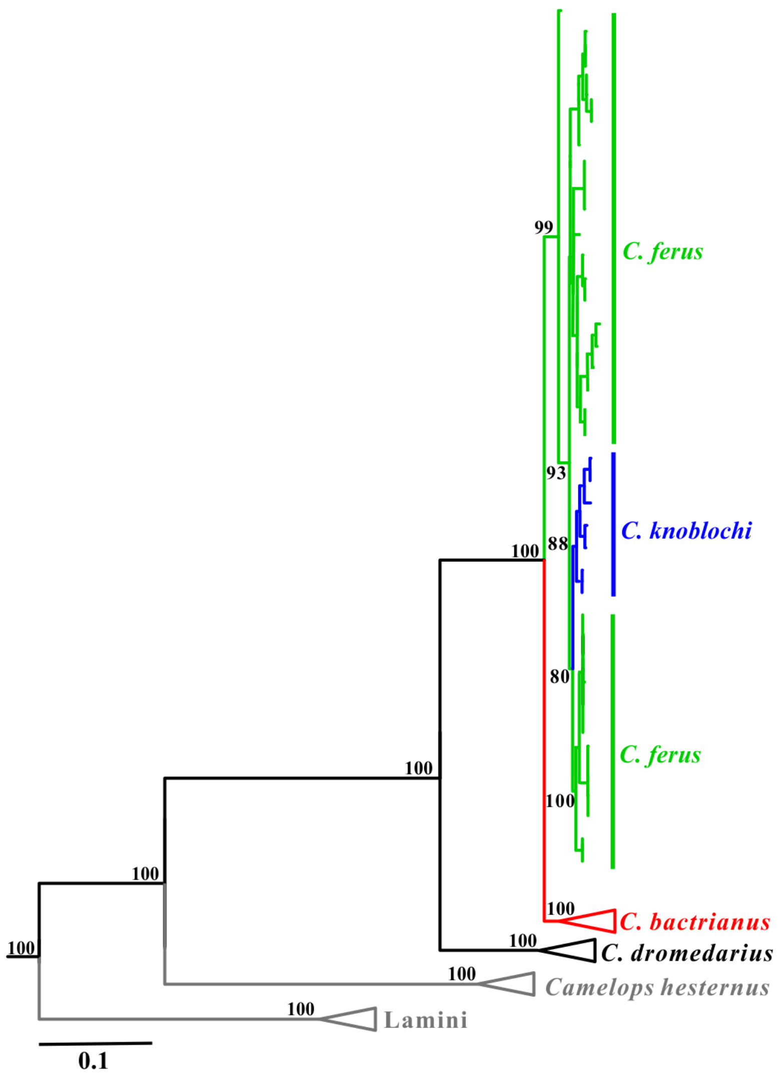


**Appendix 2-figure 4. Maximum-likelihood phylogenetic tree of camelids based on complete mitochondrial genomes**. Rooted using *Sus scrofa* as outgroup (not shown here). Branch labels show bootstrap values derived from 1,000 replicates.

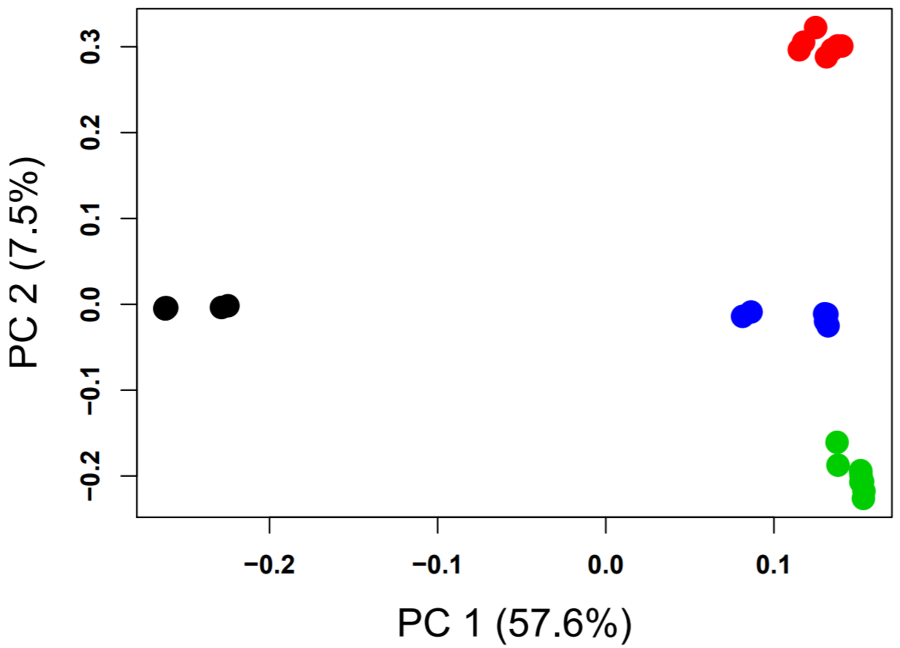


**Appendix 2-figure 5. Principal component analysis computed using genotype likelihoods with *C. dromedarius*.** The dataset used to compute these analyses contained six individuals (two from each modern species) that had aDNA damage simulated onto the reads and were downsampled to ~0.07x. Colours represent each species: black - *C. dromedarius*, red - *C. bactrianus*, green - *C. ferus*, and blue - *C. knoblochi.*

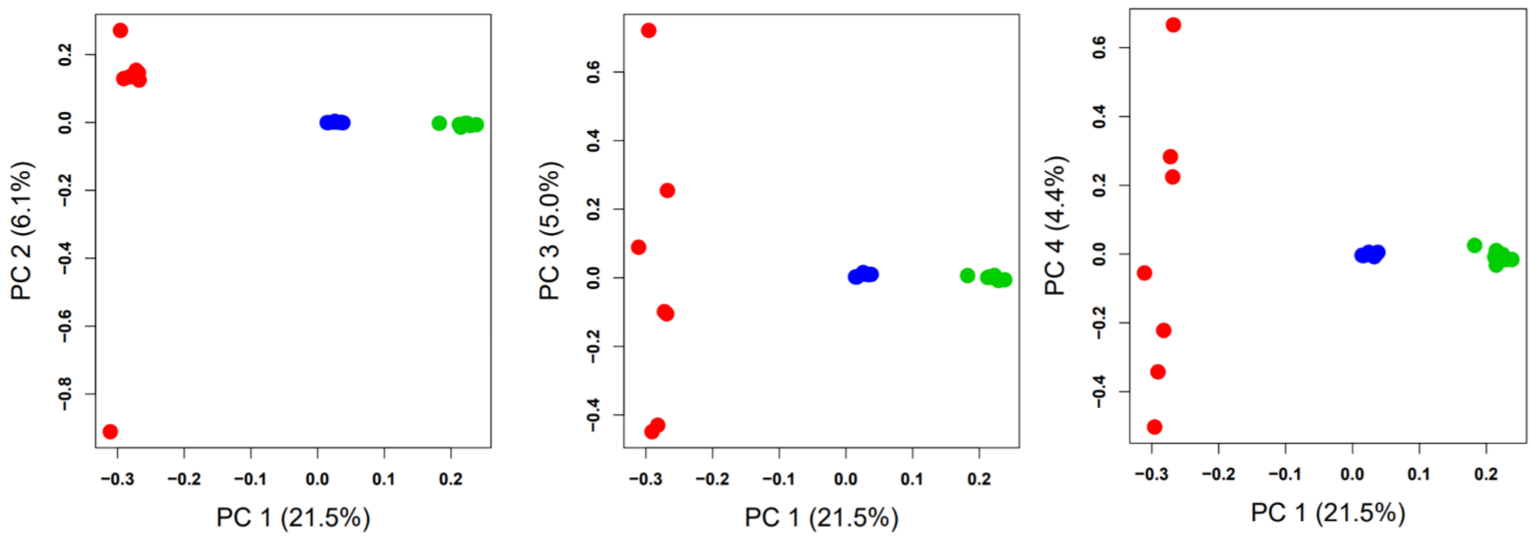


**Appendix 2-figure 6. Principal component analysis computed using genotype likelihoods without *C. dromedarius*.** Each plot shows a different combination of PC1 and either PC2, PC3, or PC4. Colours represent each species: red - *C. bactrianus*, green - *C. ferus*, and blue - *C. knoblochi.*

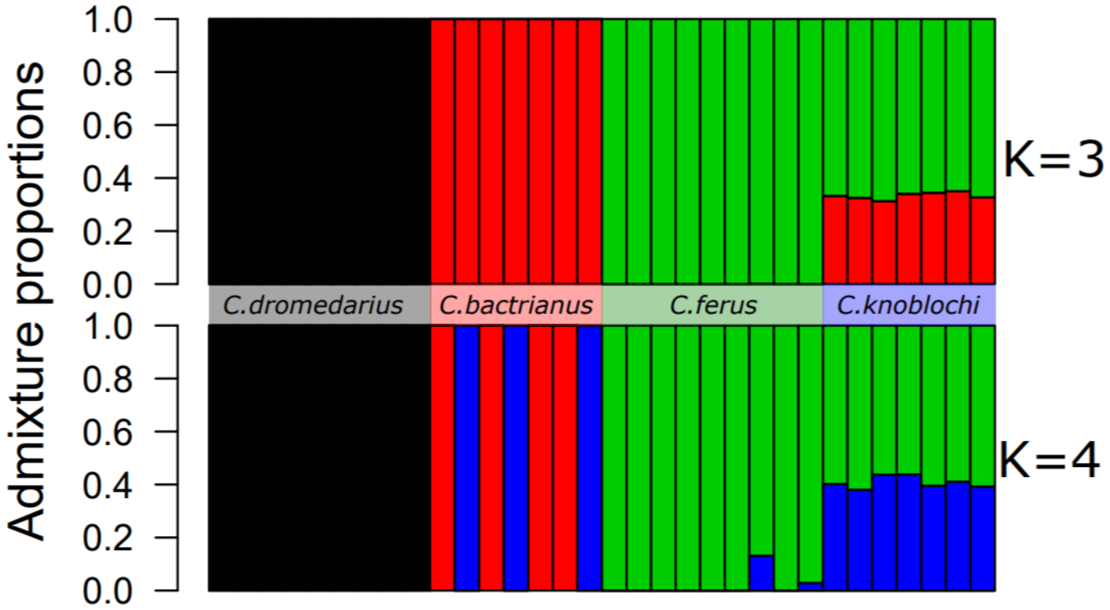


**Appendix 2-figure 7. Admixture proportions analyses computed using genotype likelihoods and an *a priori* population numbers (K) of three and four.** The dataset used to compute these analyses contained six individuals (two from each modern species) that had aDNA damage simulated onto the reads and were down sampled to ~0.07×.

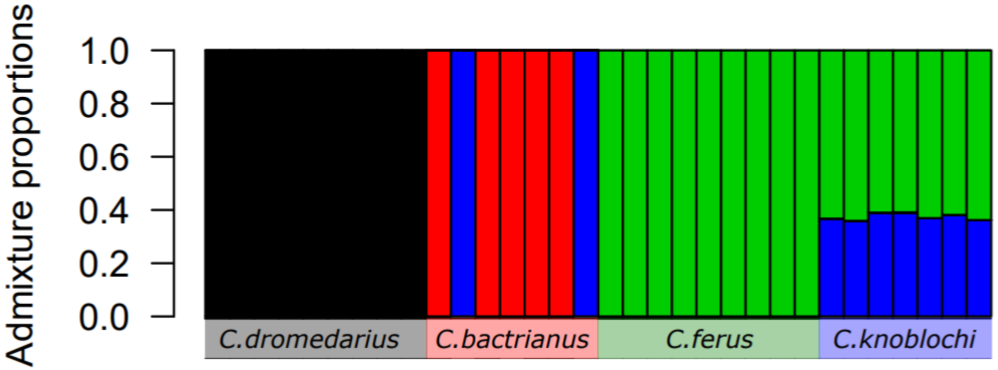


**Appendix 2-figure 8. Admixture proportions analysis computed using genotype likelihoods and an *a priori* population number (K) of four with the complete dataset.** This result had the best likelihood score but did not reach convergence.

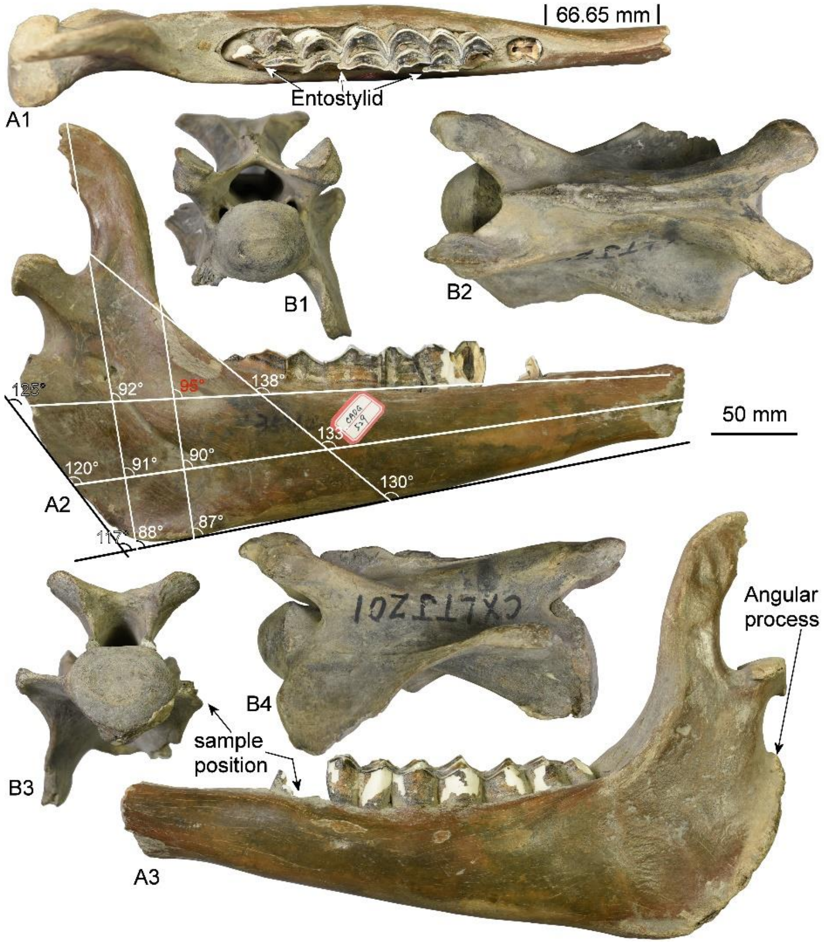


**Appendix 2-figure 9. The axial skeleton of *C. knoblochi* from Northeastern China.** A, left mandible from Qinggang, CADG529: A1, occlusal view; A2, lingual view; A3, labial view. B, the fourth cervical vertebra from Qinggang, CADG387: B1, anterior view; B2, dorsal view; B3, posterior view; B4, left lateral view. About 40% of the natural size.

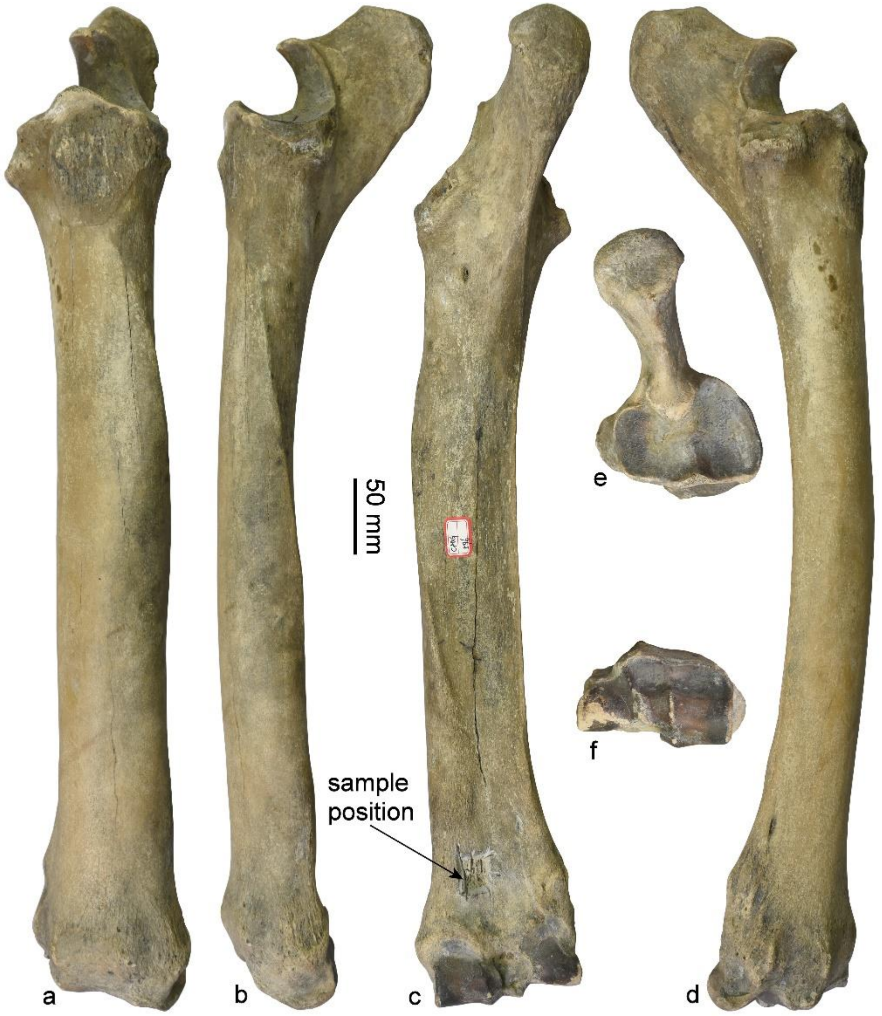


**Appendix 2-figure 10. The complete forearm of *C. knoblochi* from Northeastern China.** Right radioulnare from Qinggang, CADG596: a, cranial view; b, medial view; c, caudal view; d, lateral view; e, proximal view; f, distal view. About 30% of the natural size.

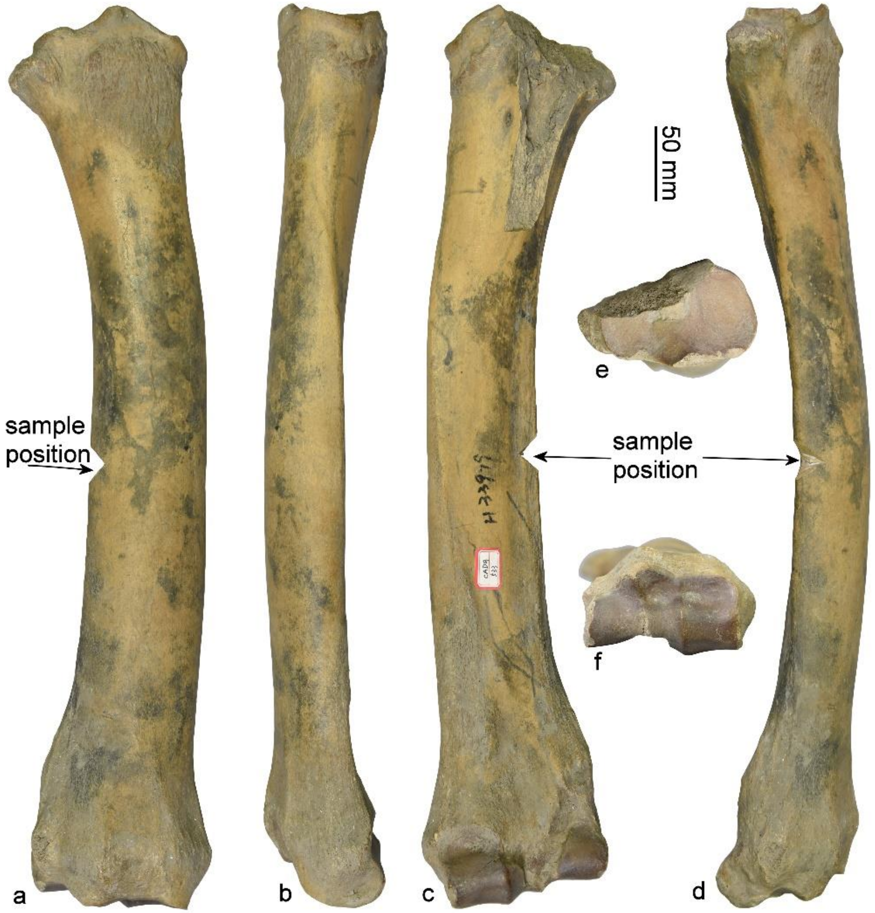


**Appendix 2-figure 11. The incomplete forearm of *C. knoblochi* from Northeastern China.** Right radioulnare from Zhaodong, CADG533: a, cranial view; b, medial view; c, caudal view; d, lateral view; e, proximal view; f, distal view. 30% of the natural size. About 30% of the natural size.

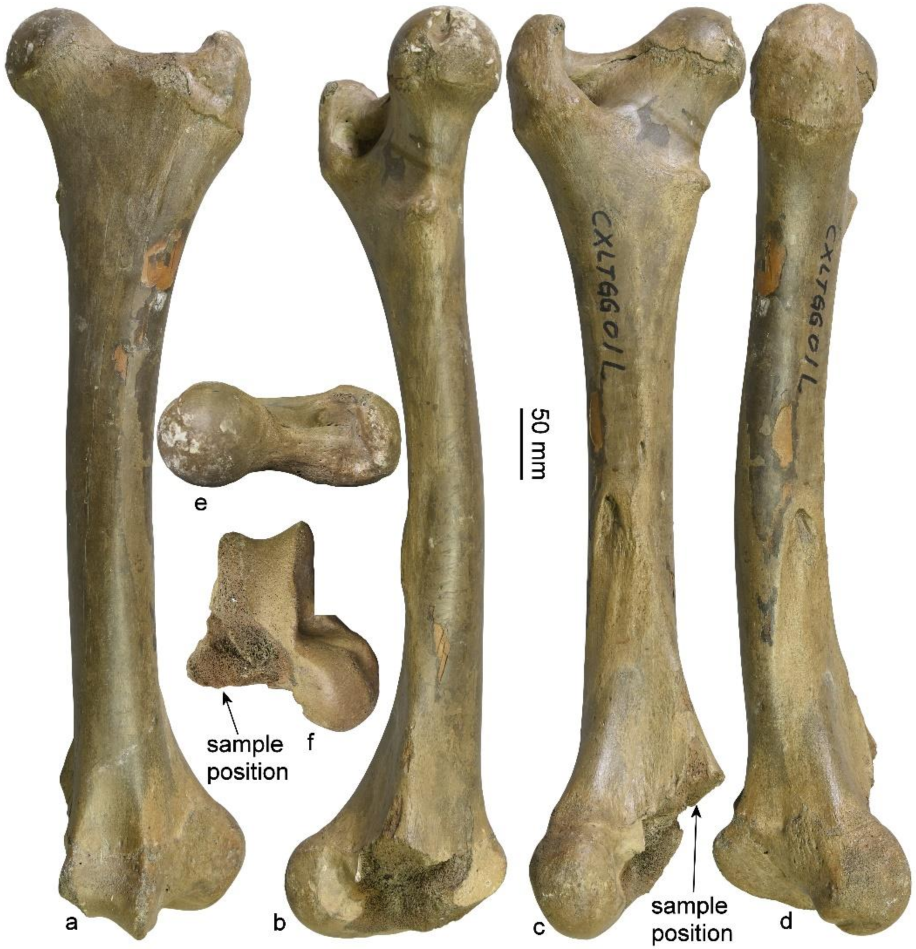


**Appendix 2-figure 12. The incomplete femur of *C. knoblochi* from Northeastern China.** Left femur from Haerbin, CADG389: a, cranial view; b, medial view; c, caudal view; d, lateral view; e, proximal view; f, distal view. 30% of the natural size.

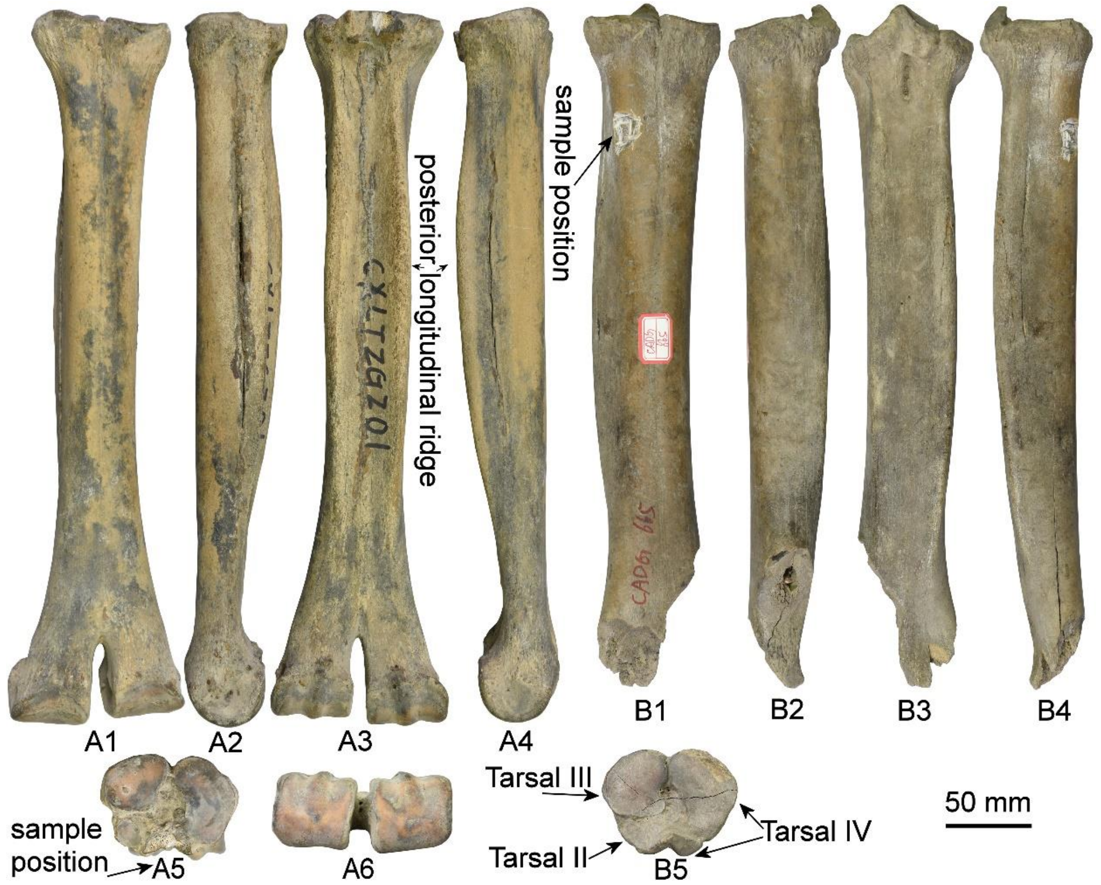


**Appendix 2-figure 13. The metatarsus of *C.knoblochi* from Northeastern China.** A, right metatarsal from Qinggang, CADG388 (nearly complete): A1, cranial view; A2, medial view; A3, caudal view; A4, lateral view; A5, proximal view; A6, distal view. B, right metatarsal from Qinggang, CADG665 (broken): B1, cranial view; B2, medial view; B3, caudal view; B4, lateral view; B5, proximal view. About 30% of the natural size.
